# Retention, not flux: endpoint confounding caps computational prediction of peptide skin penetration, with a delivery-aware reframing

**DOI:** 10.64898/2026.06.25.734657

**Authors:** Nikolaos Komianos, Pranadarth Prakash

## Abstract

Bioactive peptides are now central to cosmetic and dermatological actives, yet predicting whether a given sequence will reach its site of action in skin remains unsolved. We contend that the dominant framing, predicting a single binary “skin permeability” label from sequence, is ill-posed, and that this, rather than a shortage of modelling power, explains the field’s stalled predictive performance. The scope of the claim is narrow: barrier-*crossing* propensity is a legitimate, learnable function of molecular structure, whereas the vehicle- and endpoint-agnostic binary label that the literature supplies is not. We support this with a first-principles analysis and a study of public-source data. First, the experimental endpoint most commonly reported, transdermal flux into a diffusion-cell receptor compartment (OECD Test Guideline 428), conflates two opposite outcomes (genuine deep delivery and undesired systemic transport) and is, for a cosmetic active, frequently a *failure* signal rather than a success signal. That receptor flux is an imperfect measure of cutaneous bioavailability is long established in dermatopharmacokinetics; our contribution is to show that the same confound, inherited through scraped labels, is what caps machine learning from sequence. Second, reported “permeability” is a property of the sequence *×* delivery-vehicle *×* measurement-compartment triad, two terms of which are usually unrecorded. Third, on public-source data, a physicochemical intrinsic-permeability estimate (Potts–Guy) carries no positive predictive signal for scraped penetration labels (grouped AUC 0.45, 95% CI 0.40–0.51); sequence-only classifiers plateau in the mid-0.70s with diminishing returns as labels accumulate (AUC 0.70–0.77); and the same descriptor pipeline on a clean single-endpoint membrane dataset scores materially higher (AUC 0.83, non-overlapping CI). Our proposed reframing separates barrier-crossing (data-driven, sequence-level) from depth-and-retention (physics-driven, delivery-aware) and treats intrinsic transdermal flux as a regulatory *risk* axis; we close by proposing a triad-annotated reporting schema and a seed benchmark.

## 1 Introduction

Topically applied peptides, including signal peptides such as palmitoyl-pentapeptide-4, neurotransmitter-inhibiting peptides such as acetyl-hexapeptide-8, and carrier peptides such as the copper tripeptide GHK, anchor a large and growing fraction of the cosmetic-actives market [18]. Their efficacy depends on reaching a specific skin stratum: the viable epidermis (keratinocytes, melanocytes) or the dermis (fibroblasts, extracellular matrix). Whether a peptide reaches that target is difficult to predict, and the cost of empirical screening (synthesis, formulation, *ex vivo* diffusion-cell assays) is high.

This has motivated repeated attempts to predict “skin permeability” from peptide sequence. Such efforts inherit a binary framing, permeable versus not permeable, and the hope that, with enough labelled examples, a model will learn the mapping. We contend that this specific framing is the problem. The barrier itself, the dependence on delivery vehicle, and above all the ambiguity of the measured endpoint combine to make the naive “permeability” label a quantity that is not well defined as a function of sequence, and therefore not learnable as a single sequence-labelled binary, even though the underlying barrier-crossing propensity is. The distinction matters. We do not claim that sequence carries no information about skin transport, which it plainly does; we claim that the label the field trains on fuses three variables and two endpoints into one target, and that no amount of that label breaks the resulting ceiling.

## 2 The barrier makes bare peptides a worst case

The stratum corneum (SC) is a ~10–20 *µ*m “brick-and-mortar” layer of keratinised corneocytes embedded in highly ordered intercellular lipid lamellae. Passive permeation is gated by partitioning into, and diffusion across, this lipid phase. The widely cited “500 Dalton rule” [1] captures the empirical observation that molecules above ~500 Da rarely achieve appreciable passive flux. The Potts–Guy relationship [2], log *K*_*p*_ (cm h^−1^) = 0.71 log *P* − 0.0061 *· M*_*w*_ − 2.72, formalises the joint penalty of size and the requirement for balanced (log *P* ≈ 1–3) lipophilicity.

Peptides are penalised on all three axes simultaneously. Most cosmetic peptides exceed 500 Da; the peptide backbone is an extended hydrogen-bonding ladder, incurring a high desolvation cost on entering the lipid [3]; and many carry net charge at skin-surface pH, incurring a large electrostatic (Born) penalty on partitioning into a low-dielectric medium [4]. Bare peptides are thus close to the worst case for passive skin transport. Consistent with this, transdermal “success” in the literature is predominantly formulation-conferred, via chemical penetration enhancers, lipidation (palmitoylation, myristoylation), encapsulation (liposomes, solid lipid nanoparticles), or physical methods (microneedles, iontophoresis). The bare sequence is rarely the active agent of penetration.

## 3 The endpoint confound: receptor flux versus tissue retention

The deeper problem is the measured endpoint. The standard *in vitro* model, a Franz diffusion cell with excised or reconstructed skin over a receptor chamber as codified in OECD Test Guideline 428 [15], has no microcirculation. *In vivo*, a molecule reaching the dermis is swept into systemic circulation; *in vitro*, with no clearance, any molecule that reaches the deep dermis eventually partitions down into the receptor fluid. Receptor-compartment appearance is therefore reported as proof of barrier breach and used as a proxy for “delivered deep enough.”

This conflates two opposite outcomes:

1. *Genuine localised delivery*: the active reaches the viable epidermis or dermis and is retained there to act on its target, which is the cosmetic goal.
2. *Through-transport*: the active crosses all layers and exits into the receptor (*in vivo*, the bloodstream). For a cosmetic this is not success; it is the regulatory boundary at which a product behaves like a transdermal drug.

For a cosmetic active, high and rapid receptor flux is frequently a failure mode rather than a success. The desirable profile is high tissue retention in the viable epidermis or dermis with low receptor flux. Retention and onward flux are, moreover, governed by partly opposing physicochemistry. A peptide that binds tissue (cationic peptides associate with anionic keratin; lipophilic peptides partition into membranes; large peptides diffuse slowly) forms a depot, with high retention and impeded onward flux, whereas a small, inert, moderately lipophilic molecule fluxes through, with low retention and high systemic transfer. The two endpoints are different axes of the same process, and the literature routinely reports the wrong one for the cosmetic question.

This is not, in itself, a new observation. That receptor flux poorly represents *cutaneous* (as opposed to transdermal) bioavailability is a foundational premise of dermatopharmacokinetics: tape-stripping the stratum corneum and assaying drug as a function of depth and time was developed precisely because total flux fails to localise where a topically applied molecule resides, and it underpins topical-product bioequivalence methodology [16, 17]. Our contribution is not to rediscover the retention-versus-flux distinction but to draw out two consequences that have not been made explicit: that this confound, inherited silently through scraped labels, is the specific reason sequence-based machine learning stalls (Section 5); and that the distinction can be turned into a computational decomposition usable for *in silico* screening (Section 6).

Layer-resolved studies make the confound concrete. For palmitoyl-KTTKS (Matrixyl), tape-stripping plus tissue extraction recovers the peptide as SC ≈ 4.2, viable epidermis ≈ 2.8, dermis ≈ 0.3 *µ*g*·*cm^−2^, with none detected in the receptor [5]; this is an ideal cosmetic depot that a flux-based readout scores as zero. Unmodified KTTKS is undetectable in every layer. Acetylhexapeptide-8 (Argireline) is recovered almost entirely in the SC, with a small fraction reaching the viable epidermis and nothing in dermis or receptor [6], surface-confined and again invisible to a flux endpoint. GHK-Cu shows quantified copper retention concentrated in the stratum corneum, with lower levels in the deeper layers and measurable permeation, tracked as copper rather than as the intact tripeptide [7]. These anchors derive from animal or excised-human skin, which is generally more permeable than intact human skin in vivo, so the absolute values are indicative and do not transfer directly to intact human skin; layer-resolved data of this kind are in any case rare, and the overwhelming majority of reports give only total flux.

## 4 Permeability is a property of sequence *×* vehicle *×* compartment

Because formulation confers penetration, a single sequence yields different “permeability” labels under different vehicles, and different *meanings* under different measurement compartments. A label scraped as “peptide X penetrated skin” is therefore underdetermined: it omits the vehicle that did the work and the compartment that was measured. Training a sequence-only classifier on such labels mixes vehicles, endpoints, and assays into a single target, and the model is asked to learn a function of sequence that is in fact a function of three variables, two of which are unrecorded. The same bare sequence (for example KTTKS) is correctly labelled non-penetrant, while its palmitoylated form is labelled penetrant, and a sequence-only featurisation that ignores the modification cannot represent the difference at all.

## 5 Empirical consequences

The three claims below are demonstrated on public-source data with standard off-the-shelf descriptors, so that they depend only on the published literature, with no reliance on any proprietary system.

### Data and methods

We assembled peptide skin-penetration labels from public sources, namely open-access primary literature and granted patents, yielding 514 unique peptide species (323 labelled penetrant, 191 non-penetrant) after collapsing duplicates by majority vote. The 63% positive rate itself reflects reporting bias, since databases preferentially catalogue peptides shown to penetrate; we therefore report AUC, which is threshold-free and prevalence-robust. The negative labels are drawn from measured non-penetration (for example, peptides reported as below detection in the receptor) and from patent screening series; because published failures are scarce, the negative class, like the positive class, is a convenience sample. The modified molecule is the unit of analysis: bare KTTKS and palmitoyl-KTTKS are distinct entries, as they must be. Each peptide is described by standard physicochemical descriptors computed from sequence (molecular weight, an additive octanol log *P*, topological polar surface area, hydrogen-bond donors and acceptors, rotatable bonds, aromatic-ring count, net charge at pH 5.5, mean Kyte–Doolittle hydropathy, and amino-acid composition; lipidation, acetylation, and amidation are encoded explicitly). These additive, sequence-derived descriptors, in particular octanol log *P* and topological polar surface area, are approximate for peptides because they do not capture intramolecular hydrogen bonding or charge state; we use them deliberately as a standard off-the-shelf baseline and make no attempt to optimise the representation. Classifiers are gradient-boosted trees evaluated under sequence-grouped 5-fold cross-validation by modified-molecule identity, so that no molecule is shared between train and test; any residual leakage from near-homologous sequences in different folds would inflate the skin AUC and so cannot account for the ceiling reported below. AUCs carry 95% confidence intervals from 2,000 bootstrap resamples. As a clean-endpoint control we use the CycPeptMPDB [10] PAMPA membrane-permeability set (7,298 measurements, 7,229 unique structures; threshold log *P*_app_ ≥ −6.0), modelled with the same descriptor family and grouped by structural identity. The analyses use only public-source data and standard descriptors; they do not use Enloq’s models, features, or calibration data. Results are summarised in Figure 1.

**Figure 1:**
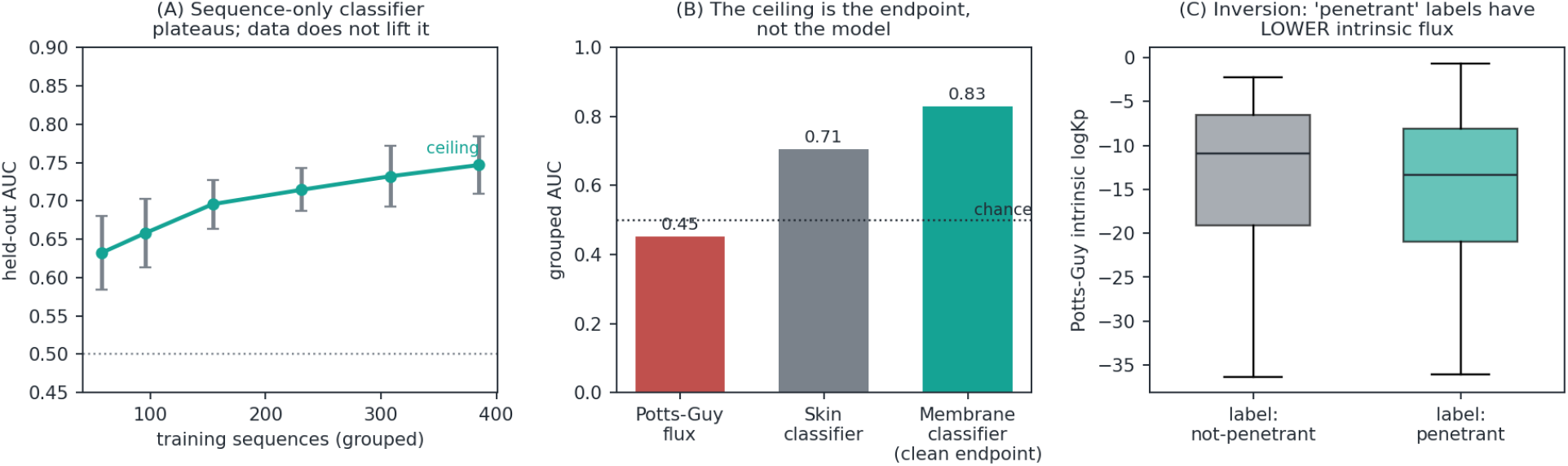
The skin-prediction ceiling is an endpoint problem, shown on public-source data. **(A)** Grouped cross-validation learning curve for a sequence-only classifier on 514 public-source peptide skin labels: AUC rises then slows in the mid-0.70s with diminishing returns (mean *±* s.d. over 24 grouped resamples; dotted line marks chance). **(B)** The same descriptor pipeline in three settings: a physicochemical Potts–Guy permeability score is at or below chance against the labels (0.45), the sequence classifier plateaus (0.70 on the shared descriptor block), and a clean single-endpoint membrane dataset (CycPeptMPDB PAMPA) reaches 0.83. The gap is consistent with the endpoint, rather than the model, as the binding constraint. **(C)** The confound made visible: peptides labelled “penetrant” have, if anything, lower estimated intrinsic permeability, indicating that the labels and the passive-permeability physics track different quantities. The analyses use only public-source data; no proprietary model is used.

#### (i) No harmonised public peptide-skin dataset of meaningful size exists

Human-skin *K*_*p*_ databases (HuskinDB [12]; SkinPiX [13]) are quantitative but predominantly *small-molecule*; peptide-permeability databases (CycPeptMPDB) measure *membrane*, not skin; topical-peptide repositories (TopicalPdb [14]) catalogue delivery without harmonised, vehicle-resolved permeation labels. Usable peptide-skin labels exist only scattered across patents and primary papers, vehicle- and endpoint-confounded as in Section 4.

#### (ii) A physicochemical intrinsic-permeability estimate carries no positive signal for the labels

The Potts–Guy intrinsic-permeability (*K*_*p*_) score, evaluated directly against the scraped penetration labels, achieves grouped AUC 0.45 (95% CI 0.40–0.51), at or below chance; because the interval includes 0.5, this establishes the absence of positive signal rather than a statistically significant inversion (Figure 1B). The evidence for the direction is distributional: peptides labelled penetrant have, if anything, *lower* estimated intrinsic permeability than those labelled nonpenetrant (Figure 1C). This is Section 3 made quantitative. A passive-permeability QSAR ranks small, lipophilic, fast through-transporters highest, while the labels reward delivery-formulated, tissue-retained peptides; the estimate correctly predicts a different endpoint from the one the labels encode. Potts–Guy was fitted on small molecules and is extrapolated well outside that domain here, which is itself consistent with its lack of positive signal on peptides.

#### (iii) Sequence-only classifiers plateau, and more data does not lift them

Under grouped cross-validation, a standard descriptor classifier reaches AUC 0.70 (95% CI 0.66–0.75) on the shared physicochemical block and 0.77 (0.73–0.81) with the richer sequence featurisation, squarely in the mid-0.70s. A learning curve built by grouped subsampling (Figure 1A) shows AUC climbing from 0.63 at ~60 training sequences to 0.75 at ~390, with diminishing returns: each doubling of labels buys progressively less, and the curve slows toward, rather than breaks through, the mid-0.70s, so the asymptote is inferred rather than reached. (The curve’s endpoint sits just below the headline 0.77 because each point is fit on a 75% grouped subsample of the training set.) Two observations bound the interpretation. First, the same descriptor pipeline on a clean single-endpoint membrane set reaches AUC 0.83 (95% CI 0.82–0.84), non-overlapping with the skin interval (Figure 1B); the membrane set differs from the skin set in size and chemistry, so this does not constitute a controlled ablation; it nonetheless shows that the descriptor family and model can discriminate materially better on a single, well-defined endpoint. Second, the diminishing returns of the skin learning curve argue that the skin–membrane gap is not closed simply by the membrane set’s larger sample size, since the skin curve is already slowing well below the membrane data volume. Taken together, these are consistent with endpoint confounding in the skin labels, rather than representational poverty of the descriptors or sheer label count, being the limiting factor. The same logic applies beyond our membrane control: cell-penetrating-peptide classifiers, which predict a single, well-defined cellular-uptake endpoint from sequence, reach substantially higher discrimination [11], whereas the skin labels here fuse retention and flux across vehicles. The published successes of peptide-permeability prediction therefore support the present thesis: they are obtained where the endpoint is unambiguous. Consistent with this, synthetic negatives generated from physicochemical boundary rules (for example Potts–Guy) do not help and can hurt, because they sharpen a trivial size and polarity axis at the expense of the genuinely informative, formulation-dependent cases.

## 6 A reframing

The remedy is to stop predicting a single confounded “permeability” and instead predict the quantities that are well posed and decision-relevant:

- *Barrier crossing*: a sequence-level probability that a peptide (as a defined molecule, modifications included) meaningfully penetrates the SC barrier. This is the data-supported question and is appropriate for machine learning.
- *Depth and retention under a given delivery system*: where in the skin a peptide is retained (SC, viable epidermis, dermis) and whether it leaks to the systemic compartment, as a function of the formulation. Because layer-resolved data are scarce, this is best addressed with a first-principles, compartmental physics model calibrated to the few quantitative anchors, rather than by data-hungry learning.
- *Intrinsic transdermal flux as a risk axis*: the physicochemical permeability estimate that fails as a success predictor is valuable as a systemic-leakage and regulatory-risk indicator, since a high intrinsic permeability, and the through-flux it implies at a given dose, flags a peptide likely to behave as a transdermal drug, which is the boundary a cosmetic must not cross.

Under this reframing, apparent paradoxes resolve. Palmitoyl-KTTKS, scored low by any flux model, is correctly a cosmetic positive once the objective is viable-epidermis retention with negligible receptor flux. Over-lipidation is correctly penalised as stratum-corneum trapping. And the recommended delivery modification for a bare sequence (for example lipidation for a moderately hydrophilic signal peptide, or an encapsulation strategy for a strongly charged one) becomes a defined, physically motivated output rather than an opaque binary.

## 7 A reporting schema and a seed benchmark

Diagnosis is cheap; the field’s bottleneck is data that is not confounded at the point of recording. We therefore propose a concrete, adoptable standard rather than a general exhortation.

### 7.1 The triad-annotated record

Every penetration datum should be reported as a structured record over the three variables that jointly determine the outcome, plus the endpoint that was actually measured (Table 1). A datum missing `vehicle, compartment`, or `endpoint` should be treated as unlabelled for modelling purposes, not as a noisy positive.

**Table 1:**
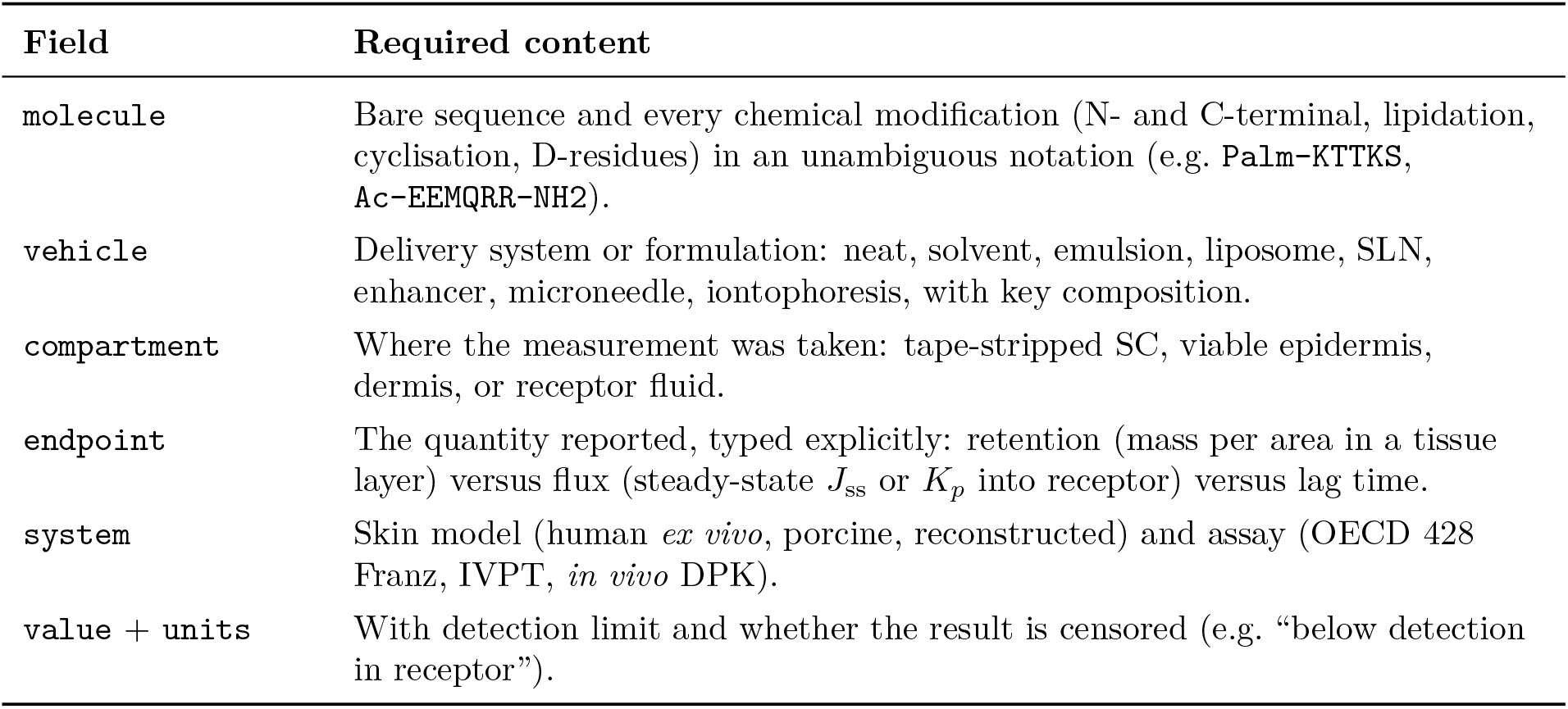
The triad-annotated record. Each penetration datum is reported over the molecule, vehicle, and compartment that jointly determine the outcome, with the measured endpoint typed explicitly.

| Field | Required content |
| --- | --- |
| molecule | Bare sequence and every chemical modification (N- and C-terminal, lipidation, cyclisation, D-residues) in an unambiguous notation (e.g. **Palm-KTTKS**, **Ac-EEMQRR-NH2**). |
| vehicle | Delivery system or formulation: neat, solvent, emulsion, liposome, SLN, enhancer, microneedle, iontophoresis, with key composition. |
| compartment | Where the measurement was taken: tape-stripped SC, viable epidermis, dermis, or receptor fluid. |
| endpoint | The quantity reported, typed explicitly: retention (mass per area in a tissue layer) versus flux (steady-state $J_{ss}$ or $K_p$ into receptor) versus lag time. |
| system | Skin model (human *ex vivo*, porcine, reconstructed) and assay (OECD 428 Franz, IVPT, *in vivo* DPK). |
| value + units | With detection limit and whether the result is censored (e.g. “below detection in receptor”). |

### 7.2 Reporting rules

1. Report the triad. Never collapse the (molecule, vehicle, compartment) tuple to a bare “permeable/not”.
2. Type the endpoint. Distinguish retention from flux explicitly; for a cosmetic active, prefer layer-resolved tissue retention (tape-stripping plus differential extraction, or imaging mass spectrometry).
3. Treat receptor flux as a regulatory readout, to be quantified and minimised, not reported as efficacy.
4. Separate the questions in curation and modelling: barrier crossing (a molecule property, learnable) versus depth and retention (a molecule *×* vehicle property, physics-constrained).

### 7.3 A seed benchmark (PepSkin-Triad)

To make the schema operational we advocate a small, openly licensed, triad-annotated benchmark with grouped (by-molecule) train/test splits and an explicit endpoint type per row, seeded by the well-characterised, layer-resolved anchors that recur in the literature (bare KTTKS versus Palm-KTTKS [5]; Argireline [6]; GHK-Cu [7]; lipidation and microneedle series [8, 9]) and grown by community contribution under the record of Section 7.1. Even a few hundred correctly annotated rows would do more for the field than thousands of confounded ones; as Figure 1 shows, the limiting factor is label quality rather than dataset size. Releasing such a slice is compatible with proprietary modelling, since a shared benchmark standardises evaluation while leaving each group’s modelling approach its own.

## 8 Conclusion

The difficulty of predicting peptide skin penetration is not primarily a modelling-capacity problem; it is an endpoint-definition problem. “Is this peptide skin-permeable?” is nearly degenerate for bare peptides and underdetermined once formulation is considered, and, as we show on public-source data, a physicochemical permeability estimate carries no positive signal against scraped labels while sequence classifiers plateau in the mid-0.70s even as a clean-endpoint control reaches far higher. Reframing the task as barrier crossing (data-driven) plus delivery-aware depth and retention (physics-driven), with intrinsic flux repurposed as a systemic-risk signal, yields questions that are both well posed and aligned with how cosmetic actives actually work. The triad-annotated schema and seed benchmark are offered to accelerate assembly of the layer-resolved, vehicle-annotated data the field actually needs.

## Author Contributions

N.K. and P.P. contributed equally to this work. Per the CRediT taxonomy, both authors share the following roles: **Conceptualization, Methodology, Software, Validation, Formal analysis, Investigation, Data curation, Visualization, Writing – original draft, Writing – review & editing**, and **Project administration**. Both authors read and approved the final manuscript.

## Ethics

This study did not involve human participants, human tissue, or animals. All analyses used previously published, public-source data.

## Use of Generative AI

Large language models were used as tools during this work. A generative-AI multimodal extraction step assisted in mining peptide measurements from patent and literature PDFs; every extracted record was gated by deterministic validation rules and subject to human review. All scientific claims, analyses, and interpretations are the authors’ own and were verified by the authors; no text, number, or citation was accepted without human verification. No generative-AI system is an author of this work.

## Data and Code Availability

The analysis in Section 5 draws on labels compiled from the public sources cited in this manuscript (open-access primary literature and granted patents) together with the public CycPeptMPDB membrane-permeability database [10]. The curated label table and the analysis code supporting Figure 1 are not released with this preprint. No proprietary Enloq model, feature, or calibration dataset was used in the analyses reported here. The public-source analysis reported here is a selfcontained demonstration of the argument that depends on no proprietary data; Enloq additionally maintains proprietary internal benchmarks and datasets that corroborate and extend these findings and are not disclosed in this preprint. This preprint is distributed under a Creative Commons Attribution 4.0 International (CC BY 4.0) licence.

